# Spatiotemporally Programmed Release of Aptamer Tethered Dual Angiogenic Growth Factors

**DOI:** 10.1101/2024.08.08.607163

**Authors:** Deepti Rana, Jeroen Rouwkema

**Affiliations:** Department of Biomechanical Engineering, Technical Medical Centre, University of Twente, 7522NB Enschede, The Netherlands

**Keywords:** Vascular endothelial growth factor, platelet-derived growth factor, aptamers, programmed release, biomaterials, tissue engineering

## Abstract

In tissue extracellular matrix (ECM), multiple growth factors (GFs) are sequestered through affinity interactions and released as needed by proteases, establishing spatial morphogen gradients in a time-controlled manner to guide cell behavior. Inspired by these ECM characteristics, we developed an “intelligent” biomaterial platform that spatially controls the combined bioavailability of multiple angiogenic GFs, specifically vascular endothelial growth factor (VEGF) and platelet-derived growth factor (PDGF-BB). Utilizing aptamer affinity interactions and complementary sequences within a GelMA matrix, our platform achieves on-demand, triggered release of individual GFs which can be “programmed” in temporally-controlled, repeatable cycles. The platform features stable incorporation of dual aptamers specific for both GFs, functional aptamer-CS molecular recognition in a 3D microenvironment with long-term stability of at least 15 days at physiological temperature, and spatially localized sequestration of individual GFs. Additionally, the system allows differential amounts of GFs to be released from the same hydrogels at different time-points, mimicking dynamic GF presentation in a 3D matrix similar to the native ECM. This flexible control over individual GF release kinetics opens new possibilities for dynamic GF presentation, with adjustable release profiles to meet the spatiotemporal needs of growing engineered tissue.

**Graphical Abstract:** 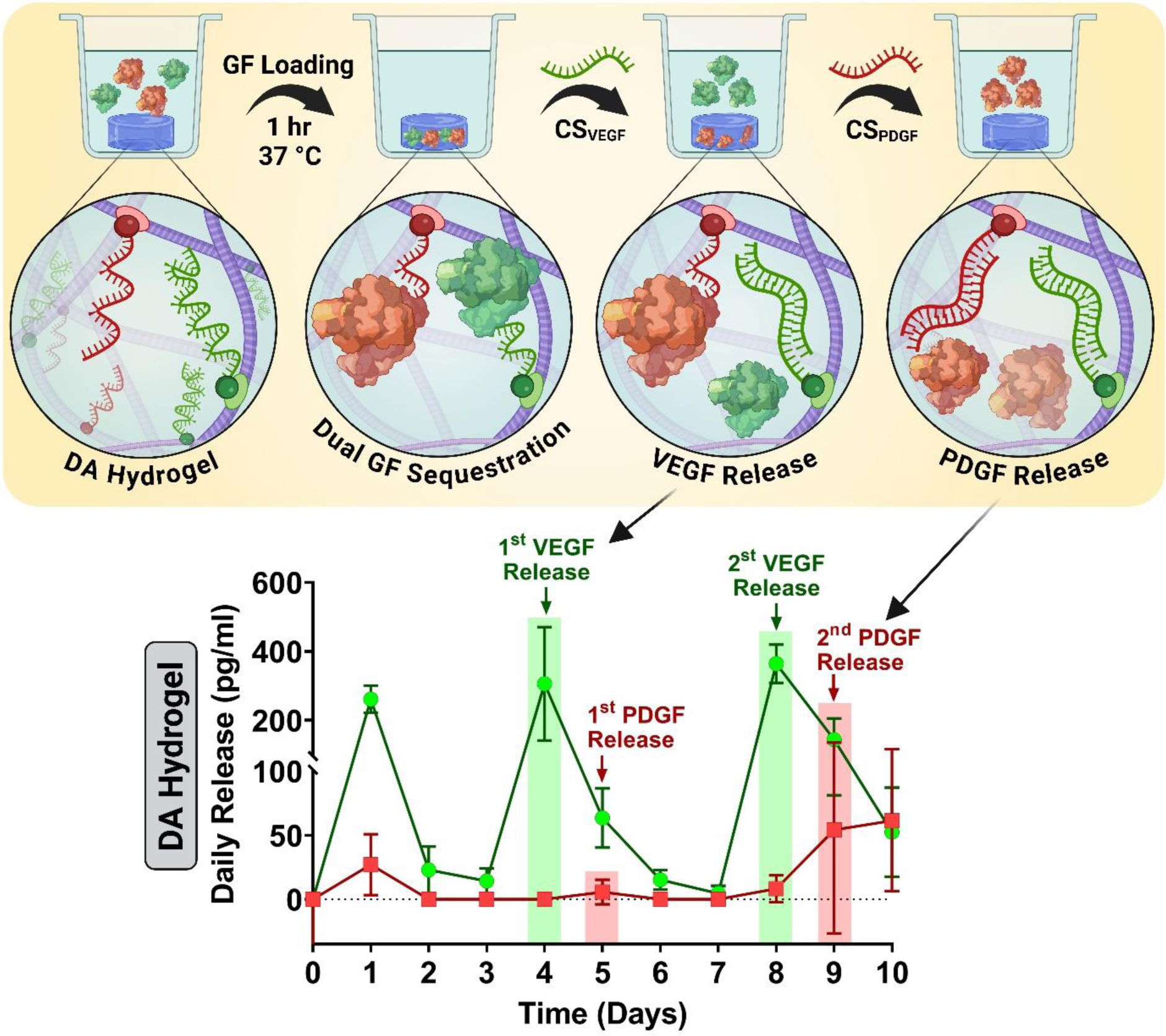

## 1. Introduction

The clinical translation of growth factor (GF) therapeutics for neovascularization of ischemic tissues and wound healing is currently hindered by their short half-life and lack of efficient delivery strategies, often necessitating the use of supraphysiological doses. For instance, FDA approved Regranex™ (becaplermin gel), developed by Ortho-McNeil Janssen Pharmaceutical (USA),[1,2] contains recombinant human platelet-derived growth factor (PDGF-BB) at 100 µg/g, which is more than 1000 times higher than physiological levels in human serum (17.5±3.1 ng/ml).[3] Such high doses, typically 7 µg/cm² applied daily for up to two weeks, are associated with significant complications, including an increased risk of cancer mortality.[4] The gel formulation results in an initial burst release, necessitating daily reapplication to maintain therapeutic efficacy. In contrast, physiological wound healing involves a temporally regulated PDGF expression, requiring targeted release over different phases. PDGF-BB’s effectiveness is also regulated by its interaction with other GFs, such as TGFβ3, which can nullify PDGF-BB’s efficacy at elevated levels.[5] This disparity between natural physiological needs and clinically available GF therapeutics highlights the need for innovative delivery strategies that can deliver multiple GFs at physiologically relevant dosages with spatiotemporal control, better mimicking the natural healing process and improving clinical outcomes.

In native tissues, multiple GFs synergistically orchestrate wound healing and neovascularization. During the early phase (0-1 day) of wound healing, platelets at the wound site express high levels of PDGF, which decreases in subsequent stages.[6] By the later phase (5-6 days), vascular endothelial growth factor (VEGF) levels, initially low, gradually peak as keratinocytes and macrophages migrate to the wound site, expressing high VEGF levels.[6] A similar temporal regulation of GFs occurs during neovascularization in newly formed granulation tissue, where high VEGF levels initially stimulate endothelial cells to form new blood vessels.[7] Later, endothelial cells express PDGF-BB, recruiting PDGFRβ receptor-expressing vascular smooth muscle cells and pericytes to stabilize and mature the vessel walls.[8] These angiogenic GFs are sequestered within the extracellular matrix (ECM) via affinity interactions and are released as needed through proteases and matrix metalloproteinases (MMPs), establishing spatial morphogen gradients that dictate cell behavior. This contrasts the traditional GF delivery systems for wound healing and tissue regeneration, which primarily rely on physical entrapment, immobilization, covalent conjugation of GFs within a polymeric matrix via degradable linkers, or microcarrier encapsulation.[9–13] While these systems achieve long-term (>30 days) sustained and sequential release profiles for dual or triple GFs both *in vitro* and *in vivo,* they typically depend on passive GF release through diffusion, polymer degradation, or proteolytic cleavage.[9,14] State-of-the-art GF delivery systems use various non-specific triggers, such as pH, light, heparin, enzymes, ultrasound, nanomaterials, cell-selective traction forces, or logic-based combinations of multiple triggers, to achieve on-demand release from biomaterials.[15–20] These methods often result in initial burst releases, polymer degradation, and sustained release upon triggering, making GFs available to tissues for extended durations. Despite their efficacy *in vitro*, these strategies have limited applicability *in vivo* due to challenges such as light scattering, lower absorption at greater tissue depths,[21] and suboptimal spatial selectivity among different GFs. Sophisticated approaches like cell-selective traction forces and MMP-based triggers address spatial selectivity issues but pose challenges such as complex system design, low cost-effectiveness, or reliance on GF modifications. Additionally, these approaches struggle with loading and releasing multiple cycles of GFs and controlling their individual release profiles. These limitations highlight the inability of current strategies to adjust GF release kinetics to meet the spatiotemporally changing requirements of the microenvironment.

Aptamer-functionalized hydrogels or microparticles have demonstrated the capability for long-term sustained release of various GFs, such as VEGF, PDGF, and basic fibroblast growth factors (FGFs), through aptamer-GF binding in both *in vitro* and *in vivo* settings.[22–24] Aptamers are small, single-stranded oligonucleotides forming unique 3D conformations that allow them to selectively bind their target GF with high affinity and specificity, minimizing the risk of non-specific release. This binding enhances GF stability, protecting them from proteolytic degradation and renal clearance, thus increasing their half-life.[25] Aptamers can also be designed to have higher affinity for their complementary sequences (CS) than the target GF, enabling specific GF release through aptamer-CS hybridization upon external triggering. However, previous studies often coupled the GF loading and hydrogel synthesis processes, exposing GF to free radicals generated during polymerization, potentially denaturing the GF and reducing their bioactivity. Not all aptamers are chemically incorporated into the hydrogel, leading to free aptamers bound to GFs, which can lower overall GF loading efficiency and result in an initial burst release.[26] These approaches primarily focused on sustaining individual GF release kinetics by exploiting aptamer affinity without exploring CS triggering[22].

Recent studies have shown promising triggered single/dual GF release profiles for up to 18 days using UV light and CS-based triggering methods.[23,27,28] However, synthesizing hydrogels with photolabile linkers involves harsh chemicals, and UV light triggering required 5 min to release GFs on demand.[27] These studies validated the bioactivity of released GFs indirectly through the supernatant. Although previous research highlights the potential for triggered GF release, upscaling these methods for 3D co-culture is challenging, as UV light exposure for as little as 30 secs can cause DNA fragmentation and cellular damage,[29] and UV light has limited penetration in deep tissues, restricting it’s *in vivo* application.[21] Additionally, while effective CS triggering has been demonstrated, the hydrogel crosslinking chemistry and GF loading methods were incompatible with cell-based studies.[23,28]

To fully explore the potential of aptamers for controlling multiple GF release kinetics, we utilized aptamers for the rapid sequestration of two angiogenic GFs (VEGF and PDGF-BB) into gelatin methacryloyl (GelMA)-based hydrogels with localized spatial control and temporally controlled individual triggered release profiles in physiological conditions. We triggered the hydrogels with CS on-demand in repeatable cycles, showcasing the “programmable” nature of GF release with control over individual release kinetics. The spatial localization and temporal release of both GFs from this platform underscore its potential for vascularized tissue engineering. We demonstrated that this biomimetic “intelligent” GF delivery platform could spatiotemporally control multiple biochemical cues, thus better mimicking the complex release kinetics of the native ECM.

## 2. Materials and Methods

### 2.1 Materials

Type A 300 bloom porcine skin gelatin (G1890-500G, Sigma Aldrich), methacrylic anhydride (MA, 276685-500ML, Sigma Aldrich), Dulbecco’s phosphate buffered saline (DPBS, D8537-500 ML, Sigma Aldrich), Fisherbrand^TM^ regenerated cellulose dialysis tubing (12-14 kDa, 21-152-14, Fisher Scientific), bovine serum albumin (BSA, A9418, Sigma Aldrich), tris(2,2‘-bipyridyl)dichloro-ruthenium (II) hexahydrate (224758, Sigma), sodium persulfate (S6172, Sigma), VEGF-specific 5′acrydite-modified aptamer (Aptamer_VEGF_, 47-nt, DNA, IDT), Aptamer_VEGF_ specific complementary sequence (CS_VEGF_, 48-nt, DNA, IDT), 5′Alexa Fluor 488-modified CS_VEGF_ (CS_VEGF-Fluoro_, 48-nt, DNA, IDT), PDGF-specific 5′acrydite-modified aptamer (Aptamer_PDGF_, 46-nt, DNA, IDT), Aptamer_PDGF_ specific complementary sequence (CS_PDGF_, 46-nt, DNA, IDT), 5′Alexa Fluor 647-modified CS_PDGF_ (CS_PDGF-Fluoro_, 46-nt, DNA, IDT), nuclease free water (11-04-02-01, IDT), 3-(trimethoxysilyl)propyl methacrylate (TMSPMA, 440159, Sigma-Aldrich), photomask films (custom-made, Selba S.A.), Fluoro-Max dyed blue aqueous fluorescent particles (2µm diameter, B0200, Thermo Scientific), Recombinant Alexa Fluor® 647 Anti-VEGFA antibody [EP1176Y] (ab206887, Lot. GR3390195-2, Abcam), Alexa Fluor 488 Anti-PDGFRα + PDGFRβ antibody (Y92, ab196376, Lot. GR3339437-3, Abcam), human VEGF-A ELISA kit (BMS277-2, Lot. 276933-006 Invitrogen), human PDGF-BB ELISA kit (RAB0397-1KT, Sigma Aldrich), glutaraldehyde solution (340855, Sigma Aldrich), vascular endothelial growth factor 165 human (VEGF, H9166, Lot. #MKCK1040, Sigma Aldrich), human PDGF-BB (P3201-10UG, Lot. #SLCC2524, Sigma-Aldrich), PDMS silicone elastomer (2401673921, Sylgard),, Corning® 96 well black polystyrene microplate (CLS3603, Sigma), Corning® Costar® ultra-low attachment 24-well plates (CLS3473, Sigma).

### 2.2 Prediction of secondary structures

The secondary structures of DNA-based aptamers and their complementary sequences were generated using NUPACK. The structure exhibiting the lowest free energy at 37°C was assumed to be the predominant conformation (Figure 1A).

**Figure 1.**
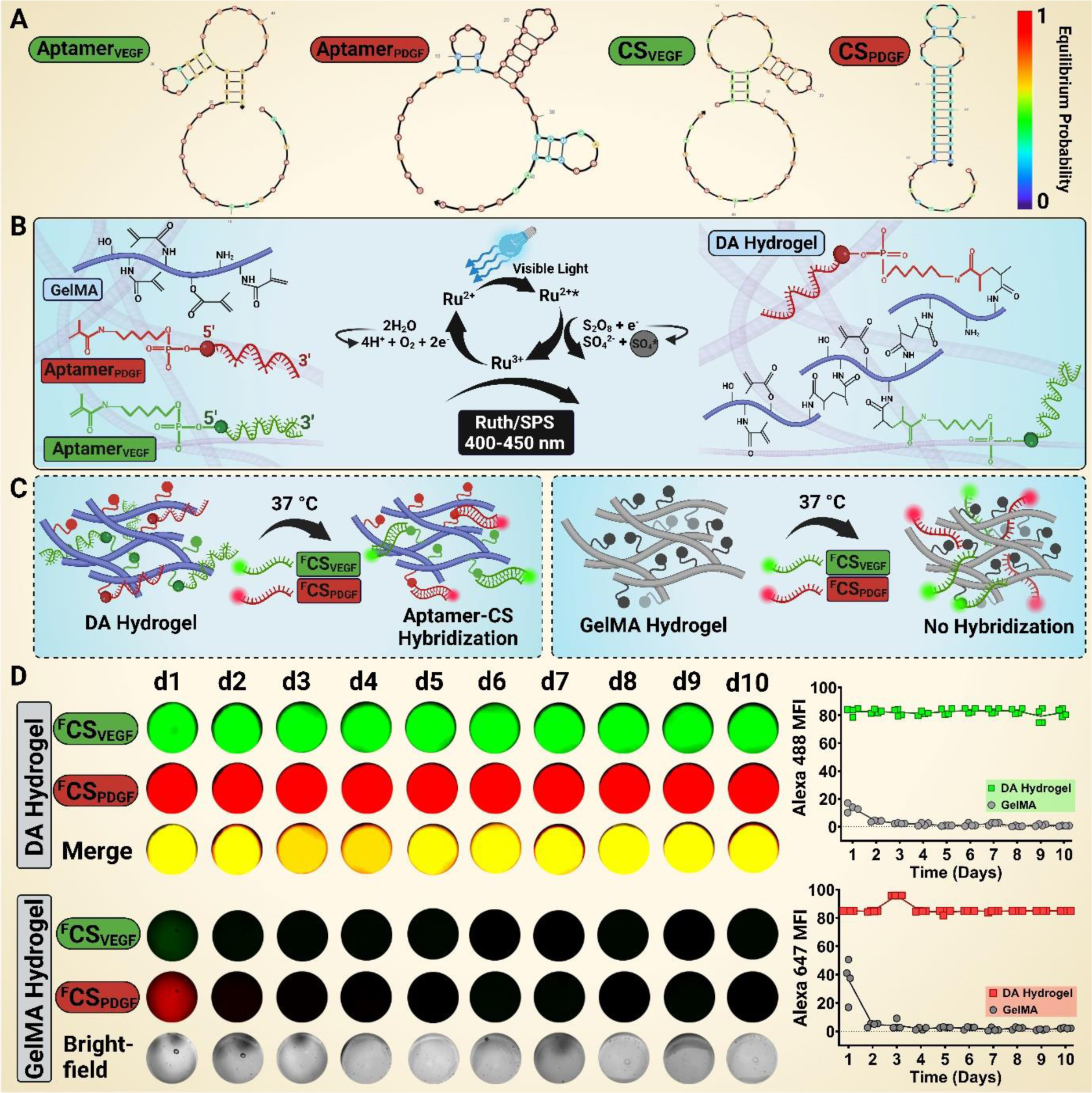
Concept of dual aptamer-functionalized (DA) hydrogels and their CS molecular-recognition. (A) Predicted secondary structures with equilibrium probability shading for the oligonucleotide sequences Aptamer_VEGF_, Aptamer_PDGF_, CS_VEGF_ and CS_PDGF_. (B) Schematic representation of photocrosslinked GelMA hydrogels containing VEGF_165_-specific Aptamer_VEGF_ and PDGF-BB-specific Aptamer_PDGF_, facilitated by ruthenium/sodium persulfate (Ruth/SPS) and visible light. (C) Diagram illustrating the concept of aptamer retention and aptamer-CS hybridization experiment. In the presence of fluorescently labelled CS (Alexa Fluor 488-modified ^F^CS_VEGF_ and Alexa Fluor 647-modified ^F^CS_PDGF_), aptamers specifically sequester their corresponding CS into the hydrogel matrix for aptamer-CS hybridization. (D) Representative fluorescence microscopic images of DA and GelMA hydrogels after 24hrs of incubation with ^F^CS_VEGF_ and ^F^CS_PDGF_ at different time-points. Brightfield images of GelMA hydrogels at all time-points are included for reference (bottom panel). The mean fluorescence intensity (MFI) of Alexa Fluor 488 and 647 in DA and GelMA hydrogels until d10. The experiment was performed with four experimental replicates (n=4), and the values are represented as the mean (connecting line) with individual data points.

### 2.3 Synthesis of dual aptamer-functionalized hydrogel

Firstly, GelMA was synthesized as previously described, achieving a medium degree of methacryloyl substitution (∼60%).[24,30] Briefly, MA (1.25% *v/v*) was added to a gelatin solution (10% *w/v*) at a rate of 0.5 mL/min for 1hr at 50 °C. The reaction mixture was then diluted five-fold with DPBS (40 °C) and dialyzed for one week using 12-14 kDa dialysis tubing (40 °C) to remove residual salts. The solution was freeze-dried for one week and stored at −20 °C.

The freeze-dried GelMA macromer (10% *w/v*) was dissolved in PBS at 50 °C for 1hr using a water-bath sonicator. Subsequently, the Aptamer_VEGF_ and Aptamer_PDGF_ were reconstituted in nuclease-free water. To fabricate <u>D</u>ual <u>A</u>ptamer-functionalized (DA) hydrogels, the pre-polymer solution was prepared with a final GelMA concentration of 5% *w/v*, Aptamer_VEGF_ at 25 µM, Aptamer_PDGF_ at 25 µM, and visible-light photoinitiators tris(2,2’-bipyridyl)dichloro-ruthenium(II) hexahydrate and sodium persulfate (Ru/SPS, 1/10 mM) in every 100 µl of pre-polymer solution. This pre-polymer solution was then used to prepare bulk DA hydrogel samples for subsequent experiments. As a control, plain GelMA hydrogels were synthesized using the same pre-polymer solution but without the aptamers. For photo-crosslinking, 100 µl of the pre-polymer solution was added into a PDMS mold, covered with a coverslip, and exposed to visible light (400-500nm, 50 mW/cm^2^) for 1 min. Samples were detached from the coverslips, transferred to ultra-low attachment 24-well plates using sterilized spatulas, and used for subsequent experiments.

### 2.4 Dual aptamer micropatterns

A two-step photopatterning method was employed to fabricate dual aptamer micropatterns. TMSPMA-treated glass slides were prepared according to a previously described protocol,[30] which introduced terminal acrylate functional groups on the glass surface, followed by sterilization via autoclaving. To pattern the Aptamer_VEGF_ region, the 1^st^ pre-polymer solution, consisting of GelMA (100 µl, 5% *w/v*), Aptamer_VEGF_ (25 µM), and Ru/SPS (1/10 mM), was added between 2mm spacers on a petri dish. This was then covered with a TMSPMA-treated glass slide and a photomask film (connected channels design) for photocrosslinking with visible light for 13s. The patterns were retrieved using warm DPBS. Subsequently, the Aptamer_PDGF_ region was overlaid on the patterned Aptamer_VEGF_ region using a 2^nd^ pre-polymer solution [GelMA (100 µl, 5% *w/v*), Aptamer_PDGF_ (25 µM) and Ru/SPS (1/10 mM)] added between 2mm spacers and photocrosslinked with visible light for 30s. The resultant micropatterns displayed Aptamer_PDGF_ regions filling the spaces between the patterned Aptamer_VEGF_ regions, allowing for the spatial localization of dual aptamers in separate regions.

For the photo-patterning of line designs for dual GF immunostaining experiments, 100 µl of the 1^st^ pre-polymer solution [GelMA(5% *w/v*), Aptamer_VEGF_ (25 µM), Aptamer_PDGF_ (25 µM) and Ru/SPS (1/10 mM)] was patterned first, followed by the 2^nd^ pre-polymer solution [GelMA(5% *w/v*), fluorescent blue microbeads (1 drop/ml) and Ru/SPS (1/10 mM)] using the same conditions as previously described. In this design, both aptamers were co-localized in one region, surrounded by a plain GelMA region.

### 2.5 Aptamer-CS molecular recognition and retention

To evaluate the retention of dual aptamers and the stability of aptamer-CS molecular recognition under physiological conditions, bulk DA hydrogels were prepared by photo-crosslinking 40 µl of DA pre-polymer solution for each sample and transferring them to a 96-well plate. These samples were incubated with Alexa Fluor-488 labelled CS_VEGF_ and Alexa Fluor-647 labelled CS_PDGF_ in 200 µl DPBS for 24hrs at 37 °C, maintaining a 1:1 mole ratio of aptamer to their corresponding CS. After incubation, the supernatant was discarded, and the samples were washed with DPBS to remove unbound CS, then supplemented with fresh DPBS (200 µl) and imaged using a fluorescence microscope (EVOS M7000, ThermoFisher Scientific). The experiment was conducted until d10, with the sample’s supernatant being replenished with fresh DPBS (200 µl) and imaged every 24hr (60% intensity, 120ms exposure time: 2x objective). Mean fluorescence intensity (MFI) was quantified using ImageJ software (NIH, USA).

To evaluate the spatially patterned dual aptamer’s CS sequestration from the media, dual aptamer micropatterns (connected channels design) were prepared as previously described. The micropatterns were transferred to ultra-low attachment well plates and incubated with both fluorescently labelled CS_VEGF_ and CS_PDGF_ in PBS (2 ml) for 24hr at 37 °C, maintain a 1:1 mole ratio of aptamer-to-CS. Subsequently, the micropatterns were washed once with PBS and supplemented with fresh PBS (2 ml). This experiment was conducted until d15, with the micropattern’s supernatant being replenished with fresh DPBS every 24hr and imaged using confocal microscopy (Nikon A1, NIS-Elements C) on d1 and d15.

### 2.6 Dual GF Immunostaining

For dual GF immunostaining, the fabricated line micropatterns were transferred to ultra-low attachment 6-well plates and incubated with a releasing medium (0.1% BSA in DPBS, 1 ml) supplemented with VEGF (50 ng) and PDGF (50 ng) for 1hr at 37 °C. The mole ratio of Aptamer_VEGF_-to-VEGF and Aptamer_PDGF_-to-PDGF was maintained at 2000:1 and 1250:1, respectively. Following incubation, the supernatant was discarded, the samples were washed once, and then incubated with releasing medium (1 ml) for 24hr at 37 °C. For GF immunostaining, the GF-loaded micropatterns were rinsed with DPBS, fixed in 4% formaldehyde solution for 30 min, and washed with DPBS. This was followed by treatment with 0.1% Triton X-100 for 10 min. The micropatterns were then washed three times with DPBS and blocked with 1% FBS for 45 min. Subsequently, the micropatterns were incubated with recombinant Alexa Fluor 647 Anti-VEGF antibody (1:100 dilution) and Alexa Fluor 488 Anti-PDGF antibody (1:50) for 48 hr at 4 °C. The immunostained micropatterns were imaged using confocal microscopy. Maximum projection images of confocal z-stacks were used to quantify the mean fluorescence intensity of VEGF and PDGF in the micropattern regions using ImageJ software.

### 2.7 Dual GFs sequestration and programmed release

To quantify programmed release behavior, bulk DA hydrogels containing Aptamer_VEGF_ (2.5 nmoles) and Aptamer_PDGF_ (2.5 nmoles) were prepared by adding 100 µl of pre-polymer solution into a PDMS mold and crosslinking it with visible-light for 1min. Plain GelMA hydrogels without aptamers were prepared as controls. After crosslinking, the hydrogels were transferred to ultra-low attachment 24-well plates. For dual GF loading, the hydrogels were incubated with 1ml of releasing medium (0.1% BSA in DPBS) containing VEGF_165_ (50 ng) and PDGF (50 ng) for 1hr at 37 °C. The mole ratio of Aptamer_VEGF_-to-VEGF and Aptamer_PDGF_-to-PDGF was maintained at 2000:1 and 1250:1, respectively. After 1hr of incubation, the supernatant was collected, and the hydrogels were washed once with releasing medium for 5min. The GF-loaded hydrogels were then incubated with 1ml of releasing medium, with the supernatant being collected and replaced with fresh releasing medium (1ml) every 24hr until d10. VEGF retention in the hydrogels was determined by subtracting the amount of free VEGF in the loading (+washing) solution after 1hr of incubation from the initially loaded VEGF amount.

The dual GF-loaded DA hydrogels were also examined for their on-demand, programmed release behavior. To trigger the first VEGF and PDGF release cycles, all hydrogel samples were incubated with CS_VEGF_ and CS_PDGF_ in releasing medium (1ml) for 24hr on d3 and d4, respectively. For the first trigger cycle, the CS-to-Aptamer mole ratios were maintained at 0.5:1 at both time-points. To trigger the second GF releasing cycles from the same hydrogel samples, all samples were incubated with CS_VEGF_ and CS_PDGF_ in releasing medium (1ml) for 24hr on d7 and d8, respectively, with CS-to-Aptamer mole ratios maintained at 2:1 for both time-points. As control, the same amount of CS was added to GF-loaded GelMA hydrogels at the same time-points. The triggered GF release was determined by the amount of GF released within 24hrs of CS addition. All supernatants, including loading and washing solutions, were stored at −20 C until further analysis.

To evaluate the efficacy of the ELISA assay in detecting GF molecules present in PBS, as a proof-of-concept, VEGF (50 ng) was added directly into 1ml of releasing medium without any hydrogel and incubated for 24hr at 37 °C. Ultra-low attachment 24-well plates were used to minimize the risk of non-specific protein adsorption. After 24hr of incubation, the solution was collected and stored at −20 °C. The amount of GF in all supernatants was measured using VEGF_165_ and PDGF ELISA kits according to manufacturer’s instructions. Absorbance for the samples was measured using a microplate reader (Infinite® 200PRO, Tecan) at 405 nm. Before analysis, supernatants were diluted with sample diluent to ensure GF concentrations were within the detectable range of the assay. Absorbance was referenced by subtracting the absorbance of zero GF concentration. ELISA data were analyzed using GraphPad Prism software, and the experiment was performed with three experimental replicates.

## 3. Results and Discussions

### 3.1 Stable dual aptamer incorporation and CS molecular recognition in DA hydrogels

We developed dual aptamer-functionalized (DA) hydrogels through photo-crosslinking using free-radical polymerization, designed for specific recognition and sequestration of VEGF_165_ and PDGF-BB. The oligonucleotide sequences, modified with acrydite groups at the 5’ end, were used to fabricate these hydrogels (Fig. 1A & Table S1, Supporting Information).[22,31,32] The hydrogel matrix was composed of GelMA, photo-initiators Ru/SPS, and the acrydite-modified aptamers (Aptamer_VEGF_ & Aptamer_PDGF_). Upon exposure to visible light at room temperature, photo-polymerization was initiated. This process allowed sulphate free radicals generated by Ru/SPS to react with the carbon-carbon double bonds of the GelMA monomers and the acrydite-modified aptamers, thus facilitating their covalent incorporation into the hydrogel network (Fig. 1B). The visible light/Ru/SPS system offers several advantages over the conventional UV light/Irgacure 2959 system, including enhanced cytocompatibility and deeper tissue penetration.[33] Additionally, Ru/SPS has reduced oxygen inhibition, enhancing hydrogel shape fidelity, and its high molar absorptivity ensures efficient crosslinking even at low concentrations.[33–35] The degree of methacrylation in GelMA is a crucial parameter influencing the crosslinking density and the successful covalent incorporation of the aptamers.[30,36,37] A higher degree of methacrylation results in more methacrylate groups available for crosslinking, thereby producing a denser polymer network that enhances aptamer incorporation by providing additional covalent bonding sites.[38] However, this increased density may also affect the spatial arrangement and stability of the incorporated aptamers, potentially influencing molecule diffusion and the accessibility of functional groups for subsequent reactions. Our previous studies have indicated that the covalent incorporation of aptamers within the GelMA network enhances the overall storage modulus of the hydrogels compared to plain GelMA hydrogels of similar methacrylation degree and monomer concentration.[24,39] Interestingly, this enhancement was independent of the aptamer amount, suggesting saturation of the available reaction sites within the GelMA network.[24] Consequently, the methacrylation degree, monomer concentration, and aptamer amount are interdependent parameters that critically determine the properties of aptamer-functionalized GelMA hydrogels. To balance bioactivity, biocompatibility,[30] and network properties, we utilized GelMA with a medium methacrylation degree (∼60%) for synthesizing DA hydrogels in this study.

GelMA, derived from gelatin, retains essential ECM characteristics, including integrin-binding motifs (RGD tripeptides) and MMP-sensitive peptide sequences, which facilitate cell adhesion and matrix remodelling.[40] GelMA also exhibits reduced antigenicity compared to collagen, tunable viscoelastic properties that regulate cell behaviors, and is a biofabrication-friendly biomaterial.[41] Its inherent GF sequestration ability, due to the net charge of gelatin chains, permits electrostatic interactions with oppositely charged GFs.[42] The hydrogel’s crosslink density, determining the mesh size of the network, modulates GF release. GFs smaller than the mesh size diffuse rapidly, whereas larger GFs are retained longer and rely on network degradation for release. To isolate the effects of aptamer incorporation on GF sequestration and release, we compared DA hydrogels with plain GelMA hydrogels of the same methacrylation degree and monomer concentration. We evaluated the retention and molecular recognition capacity of the dual aptamers within the hydrogels using 5’-Alexa Fluor 488-labelled CS specific for Aptamer_VEGF_ (^F^CS_VEGF_) and 5’-Alexa Fluor 647-labelled CS specific for Aptamer_PDGF_ (^F^CS_PDGF_) at physiological temperature (37 °C) (Fig. 1B & C).

The DA hydrogels exhibited significantly higher fluorescence for both ^F^CS_VEGF_ and ^F^CS_PDGF_ from d1 to d10, confirming stable aptamer incorporation (Fig. 1D). The mean fluorescence intensity (MFI) for ^F^CS_VEGF_ was 82.84 on d1 and 82.11 on d10, while the MFI for ^F^CS_PDGF_ was 84.98 on d1 and 85.00 on d10. In contrast, plain GelMA hydrogels showed substantially lower MFI for both CS on d1 (^F^CS_VEGF_-13.46; ^F^CS_PDGF_-36.50), which decreased significantly by d2 (^F^CS_VEGF_-4.11; ^F^CS_PDGF_-4.63) and was almost negligible by d10 (^F^CS_VEGF_-0.67; ^F^CS_PDGF_-2.12). The stable fluorescent signals in DA hydrogels throughout the study confirmed the covalent incorporation and high functionality of the aptamers for CS molecular recognition in a 3D matrix. In contrast, plain GelMA hydrogels without aptamers showed significantly lower CS adsorption and sequestration after 24 hr, with almost negligible fluorescence after 48 hr. This demonstrates the critical role of dual aptamer incorporation in enhancing the molecular recognition and retention capabilities of the hydrogels.

### 3.2 Spatially patterned dual aptamers provide localized and homogenous CS hybridization

Native tissues exhibit a complex spatial organization of multiple GFs localized in specific regions, precisely controlling cellular behavior and tissue architecture. Spatially patterned delivery of multiple GFs ensures that cells receive appropriate signals at the correct locations, promoting the formation of a structured and functional vascular network. Additionally, GF gradients guide cell migration to long-range locations, establishing zones where specific cell types dominate, such as regions rich in endothelial cells forming capillaries surrounded by supportive stromal cells. Replicating these natural GF patterns in engineered tissues can enhance functionality and integration with host tissues. Thus, developing strategies to spatially localize multiple GFs within an engineered construct, without GFs bleeding into each other’s regions, is crucial.

To assess the long-term stability of spatially localized dual aptamers within a construct, we employed a two-step photopatterning technique (Fig. 2A). First, the Aptamer_VEGF_-based pre-polymer was photo-patterned using a photomask, followed by overlaying the Aptamer_PDGF_-based pre-polymer in remaining region. To visualize the distinct patterned aptamer regions, the micropattern was incubated with both ^F^CS_VEGF_ and ^F^CS_PDGF_ for 24 hr at 37°C. The samples were then washed and incubated with DPBS until d15 at physiological temperature, with DPBS replenished every 24 hr. Fluorescence microscopy stitched images of the micropatterns on d1 and d15 revealed aptamer-specific CS sequestration within the patterned regions (Fig. 2B). The intensity profile plot across the micropatterns at both time-points displayed peaks and valleys consistent with the patterned region width for both aptamers. High fluorescence hydrogel accumulation near the patterned region interface and rough interface edges towards the Aptamer_PDGF_ regions indicated partial degradation of the Aptamer_PDGF_-modified hydrogel by d15. We previously showed that in the two-step photocrosslinking process, the regions exposed twice for crosslinking exhibit a higher mechanical modulus due to increased crosslinking density compared to regions crosslinked only once.[39] Thus, the degradation of the single photocrosslinked Aptamer_PDGF_ region likely results from changes in its crosslinking density. This disparity in network properties can be addressed by employing alternative biofabrication techniques, such as 3D bioprinting, for spatial patterning of multiple aptamers.

**Figure 2.**
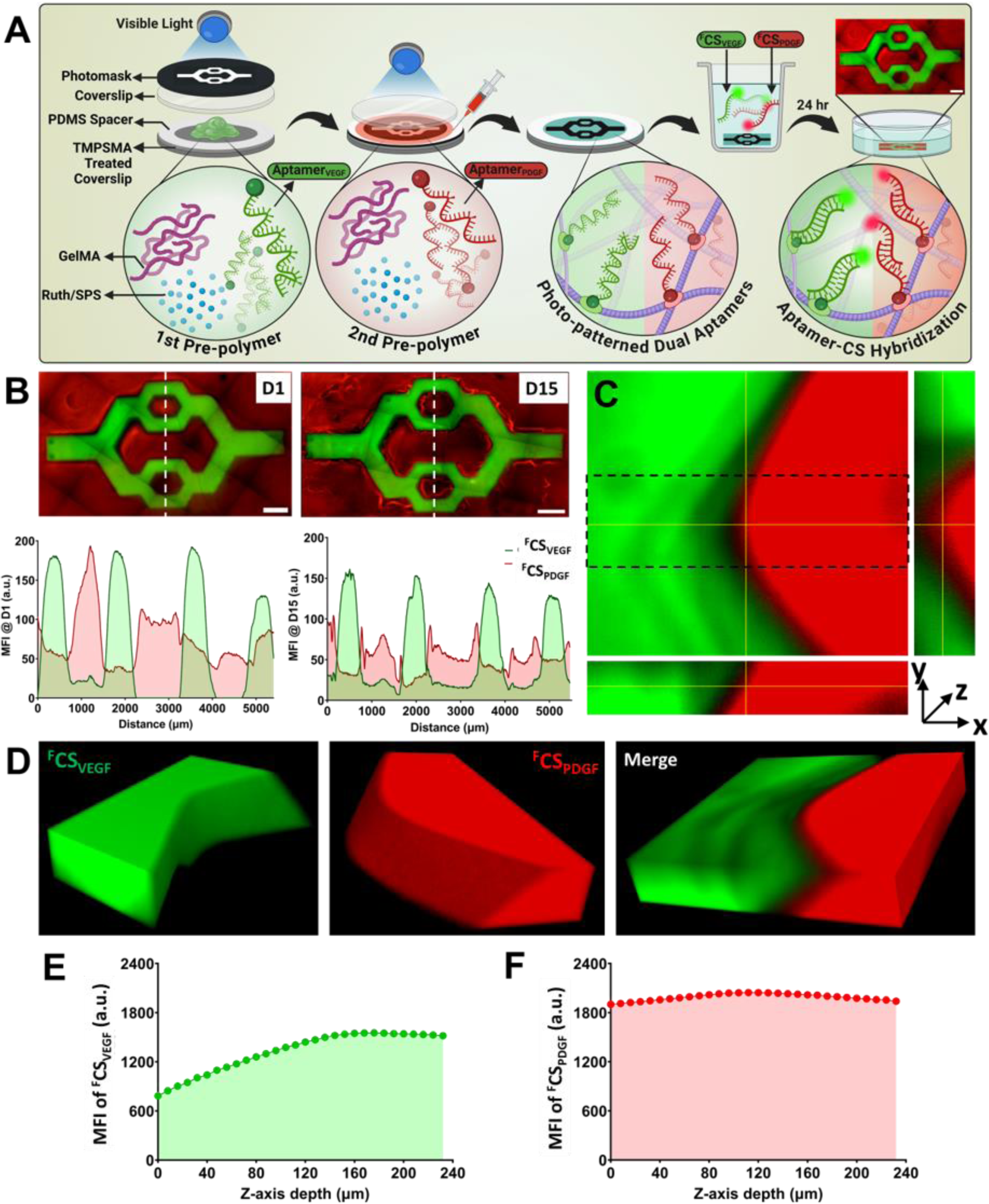
Spatially localized dual aptamer micropatterns. (A) Schematic representation of the fabrication of dual aptamer micropatterns using a two-step photocrosslinking method, followed by spatially localized sequestration of fluorescently labelled CS (^F^CS_VEGF_ and ^F^CS_PDGF_) from the culture medium after 24hrs of incubation. (B) Stitched microscopic images of the CS-sequestered micropatterns with defined Aptamer_VEGF_ (green) and Aptamer_PDGF_ (red) regions on d1 and d15 (Top). The scale bar is 1000 µm. The MFI profile of CS sequestered micropatterns as a function of distance on d1 and d15, where green and red shaded regions correspond to ^F^CS_VEGF_ and ^F^CS_PDGF_ fluorescence intensities, respectively (Bottom). The vertical white dashed line over the micropattern images at d1 and d15 indicates the measurement points for MFI profiles. (C) Orthogonal view and (D) 3D projection of confocal z-stacks of the micropattern on d1. The MF1 profiles as a function of micropattern depth along the z-axis for (E) ^F^CS_VEGF_ and (F) ^F^CS_PDGF_ fluorescence confirm homogenous CS sequestration throughout the micropattern depth. The marked region (black dashed line) in the orthogonal view of the micropattern (C) represents the region of interest (ROI) used for MFI profiles along the z-axis depth of the micropattern.

3D projections and orthogonal images (z = 240 µm) of confocal z-stacks confirmed the homogeneous sequestration of both CS throughout the patterned aptamer regions’ thickness while maintaining a clear interface between the Aptamer_VEGF_ and Aptamer_PDGF_ regions (Fig. 2C,D & Video S1, Supporting Information). The MFI of ^F^CS_VEGF_ throughout the depth of the micropattern region of interest (ROI, black dashed rectangle) showed a lower MFI at 0 µm depth, gradually increasing to 120 µm and stabilizing thereafter (Fig. 2E). The ^F^CS_PDGF_ MFI remained stable throughout the thickness (Fig. 2F). The variation in ^F^CS_VEGF_ MFI can be attributed to light deflection passing through the photomask during the photopatterning process, resulting in a patterned region with a tilted angle near the edges. These observations confirm the ability of the patterned dual aptamers to sequester specific CS into the patterned regions and retain hybridized CS within the micropatterns until d15, with no signs of aptamer/CS leakage into other regions. This demonstrates the successful spatial localization of individual aptamers and their stable aptamer-CS molecular recognition for long-term at physiological temperature.

### 3.3 Spatially localized aptamer-tethered VEGF & PDGF

To evaluate the dual GFs sequestration capability of DA hydrogels in a spatially defined manner, we fabricated line micropatterns consisting of alternating patterned DA hydrogel regions and plain GelMA regions, each with a width of 500 µm (Fig. S1, Supporting Information). This was achieved using a two-step photocrosslinking method. Blue fluorescent microbeads (diameter: 2 µm) were mixed with the GelMA pre-polymer solution to clearly demarcate the interface between the DA and GelMA regions. For GF loading, the micropatterns were incubated with both VEGF and PDGF, each at a concentration of 50 ng in 1ml of releasing medium (0.1% BSA in DPBS), for 1 hr at 37°C. Post-incubation, the micropatterns were washed with DPBS and then incubated with fresh releasing medium (1 ml) for an additional 24 hr at 37°C. To visualize the sequestered GFs, the samples were washed, fixed with paraformaldehyde, and immunostained with Alexa Fluor 647-conjugated anti-VEGF antibody and Alexa Fluor 488-conjugated anti-PDGF antibody.

Confocal z-stack imaging revealed significantly higher co-localization of VEGF (red) and PDGF (green) within the patterned DA hydrogel regions compared to the GelMA regions (blue) following 1 hr of dual GF loading (Fig. 3 & Video S2-4, Supporting Information). The 3D projections and XZ orthogonal views of the confocal z-stacks demonstrated a homogeneous and localized GF penetration throughout the thickness of the DA regions (z=240 µm) (Fig. 3A,B&D). Fluorescence intensity analysis throughout the micropattern ROI depth indicated a consistent MFI for VEGF along the z-axis, whereas PDGF intensity progressively increased from the bottom to the top layer (0 to 240 µm) (Fig. 3F). The minimal MFI of blue microbeads throughout the z-axis confirmed their limited presence within the ROI’s depth. The MFI profile plot as a function of distance across two representative regions (Fig. 3B&D) of the micropattern displayed co-localized intensity peaks for VEGF and PDGF within the DA regions, while the blue microbeads showed distinct intensity peaks in the GelMA regions (Fig. 3C&E). In line with the confocal images, normalized VEGF MFI accounted for only 1.46% in the GelMA regions, and normalized PDGF MFI was merely 0.04% in these regions.

**Figure 3.**
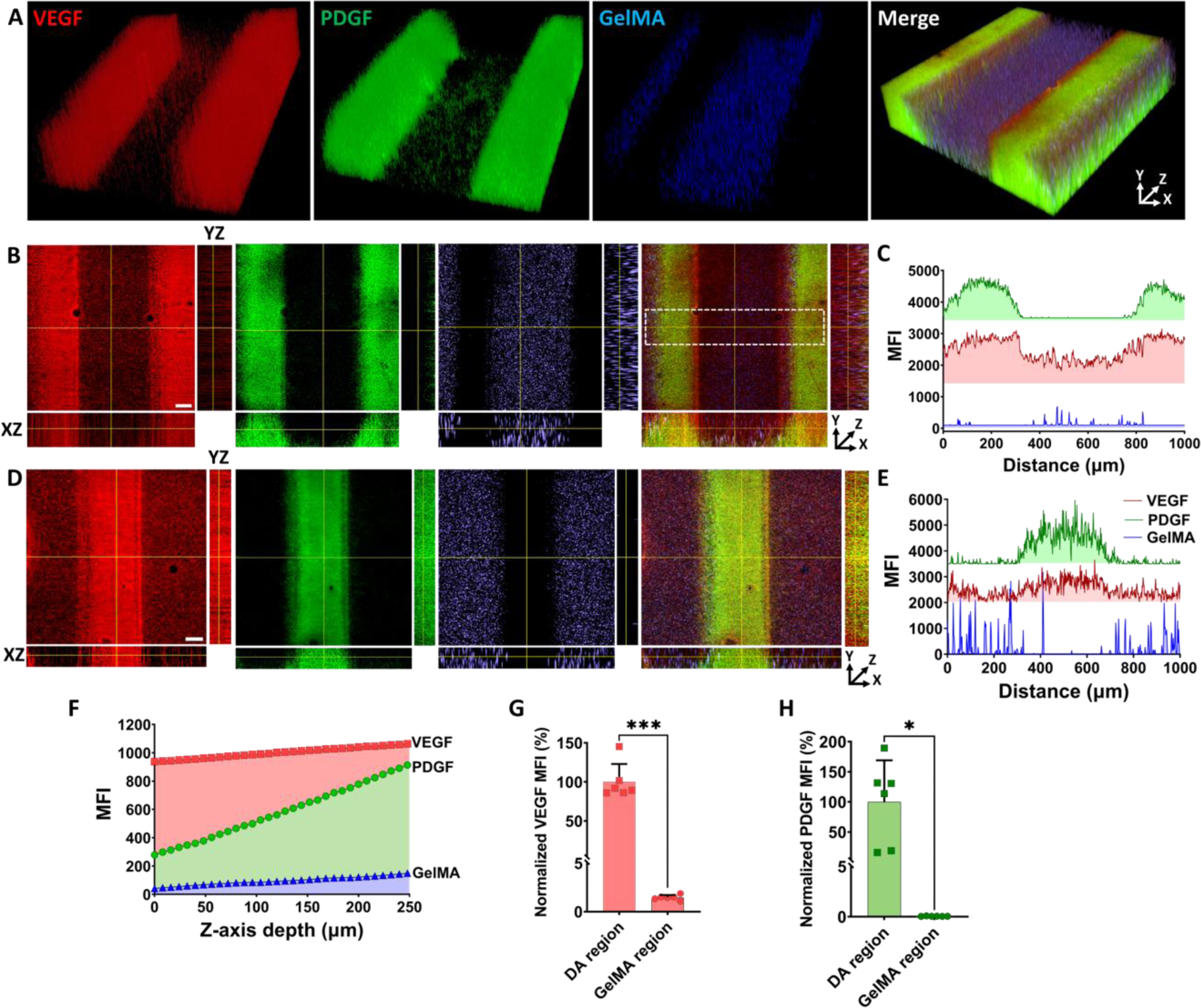
Aptamer-tethered spatial localization of VEGF and PDGF within micropatterns. (A) 3D projection of confocal z-stacks montage of VEGF (red) and PDGF (green) immunostained micropattern, confirming maximum GF sequestration within dual aptamers-patterned regions (line width-500 µm). The blue color in neighboring GelMA regions (line width-500 µm) corresponds to fluorescent microbeads mixed in the GelMA pre-polymer. (B&D) Orthogonal view montages of two different regions of the micropattern, confirming homogenous sequestration of both GFs throughout the micropattern thickness (z=240 µm). (C&E) The stacked MFI profiles as a function of distance in two different patterned regions (as indicated by yellow horizontal line in B & D) for VEGF (red), PDGF (green) and GelMA (blue) fluorescence. (F) The MFI profiles of individual fluorescence for VEGF (red), PDGF (green) and GelMA microbeads (blue) throughout the z-axis depth of the micropattern ROI (white dashed line in B). Normalized VEGF (G) and PDGF (H) MFI% in patterned DA and GelMA regions of the micropattern (n=6, technical replicates). The statistical significance was calculated using unpaired t-test with Welch’s correction where alpha was fixed at 0.05 with ***p=0.0001, and *p=0.0164.

The competitive GF sequestration observed between the DA hydrogels and the GelMA regions suggests a preferential binding of aptamers to their specific GF molecules, surpassing the gelatin’s inherent electrostatic interactions. These results, which aligns with our earlier findings on spatially localized aptamer-CS stable molecular recognition (Fig. 2), confirm successful aptamer-GF tethering. The DA hydrogels demonstrated the capability to rapidly and homogenously sequester both GFs throughout the hydrogel thickness on a macroscale within just 1 hr. The absence of GF bleeding or leakage into other patterned regions underscores the strong molecular recognition and binding specificity of the aptamer-GF interactions at physiological temperature.

### 3.4 Temporally programmed differential release of aptamer-tethered VEGF and PDGF in repeatable cycles

The precise control over the sequestration and release of GFs using aptamer-functionalized hydrogels hinges on the molecular recognition between the target GF, the aptamer, and CS oligonucleotides. Building on the demonstrated aptamer-CS hybridization and aptamer-GF tethering, we sought to quantitatively assess the dual GF sequestration from the medium and their time-controlled release via aptamer-CS hybridization. For this, photo-crosslinked DA hydrogels (100 µl/sample) containing 2.5 nmoles of Aptamer_VEGF_ and Aptamer_PDGF_ were incubated with a loading medium containing 50 ng each of VEGF and PDGF for 1 hr at 37°C (Fig. 4A & C). Post-loading, the samples were washed once with DPBS, then supplemented with fresh DPBS and incubated at 37°C until d10. The medium was replenished every 24 hr, and supernatants were collected for subsequent ELISA assays. As a proof-of-concept, the amount of VEGF sequestered after 1 hr of GF loading was quantified by analyzing the loading solution (combined with the washing supernatant for each sample) using the ELISA assay. Similar ELISA measurements were performed for PDGF sequestration in DA hydrogels. Due to the absorbance readings exceeding the ELISAsensitivity range, these sampleswere excluded. To validate the sensitivity of the ELISA assay for detecting free VEGF in the supernatant, 50 ng of VEGF was directly added to 1 ml of releasing solution (0.1% BSA in PBS), incubated for 24 hr at 37°C, and assessed using the ELISA assay (referred to as the “PBS” sample).

**Figure 4.**
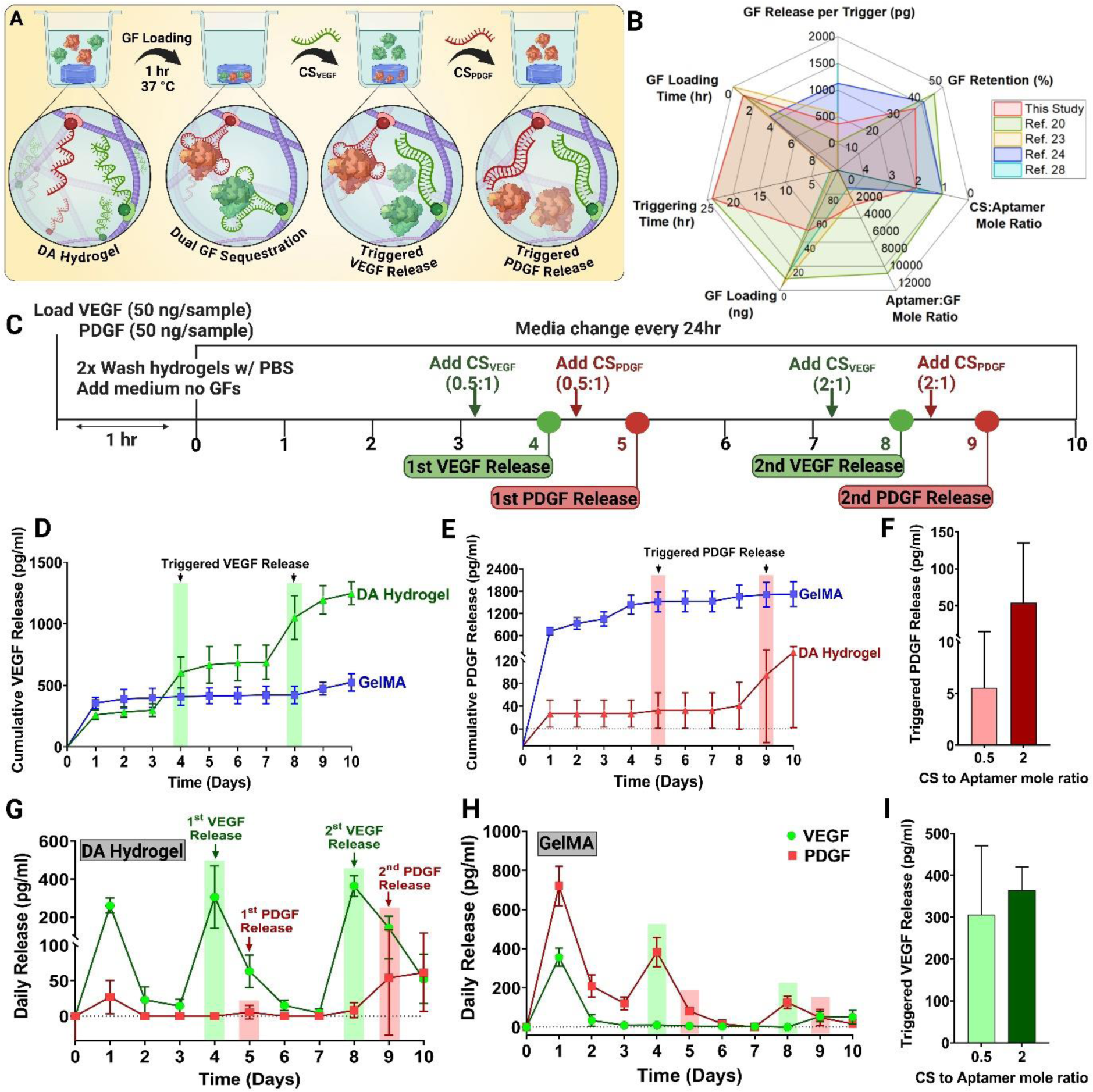
Temporally programmed release of dual GFs from DA hydrogels. (A) Illustration of the GF loading process in DA hydrogels. The release of individual GFs can be temporally controlled by adding specific CS triggers to release individual GF, independently. (B) Comparison of GF sequestration and release properties between DA hydrogels (this work) and previously reported aptamer-modified hydrogels [20,23,24,28]. For studies involving dual GF release, properties corresponding to the first GF release were considered. If specific properties of aptamer-modified hydrogels were not measured, the corresponding data are plotted in the center of the radar chart. (C) Complete timeline of the experiment. Cumulative release of (D) VEGF and (E) PDGF from DA and GelMA hydrogels over 10 days, where CS was added on specified time-points. Daily release of both VEGF and PDGF from (G) DA hydrogels and (H) GelMA hydrogels over 10 days in the presence of CS. To trigger the first and second rounds of VEGF release on d4 and d8,CS_VEGF_ was added to both hydrogels on d3 and d7 for 24hrs (as indicated in the timeline). Similarly, to trigger the first and second rounds of PDGF release on d5 and d9, respectively, CS_PDGF_ was added to both hydrogel samples on d4 and d8 for 24hrs. The CS-to-aptamer mole ratio used for the first and second rounds of release for both GFs was 0.5:1 and 1:1, respectively. (F) CS-triggered PDGF and (I) VEGF release from DA hydrogels using different CS-to-aptamer mole ratios. The experiment was performed with three experimental replicates (n=3). The graph are represented as mean ± SD.

Quantification of free VEGF, normalized to control PBS samples, revealed a higher presence of free VEGF in the loading solution of GelMA hydrogels (72.39 ± 6.78%, coefficient of variance, CV=9.37%) compared to DA hydrogels (63.37 ± 6.22, CV=9.81%) after 1 hr of loading (Fig. S2, Supporting Information). This indicates that DA hydrogels sequestered approximately 37% of VEGF from the loading solution, compared to about 28% by GelMA hydrogels within the same timeframe. The enhanced VEGF sequestration in DA hydrogels is likely attributed to the high binding affinity between the GF and Aptamer_VEGF_. Gelatin-based hydrogels, such as GelMA, naturally exhibit GF sequestration properties through electrostatic interactions with oppositely charged GFs like VEGF and PDGF. This explains the relatively high sequestration observed in GelMA hydrogels within just 1 hr of loading. Measuring the free VEGF concentration present in PBS samples using the ELISA assay showed an output of only 16.99 ± 0.59 ng/ml (CV = 3.51%), which is approximately 34% of the initially loaded VEGF concentration (Fig. S3, Supporting Information). This disparity in ELISA sensitivity towards detecting free VEGF molecules likely relates to the overall lower free VEGF presence in all samples, suggesting the actual concentrations could be higher.

To investigate the programmed release behavior of GFs, DA and GelMA hydrogels preloaded with GFs were supplemented with CS at varying mole ratios to aptamers in the release medium at specified time-points and incubated for 24 hrs. The daily release profile of DA hydrogels demonstrated a triggered release of individual GFs from the same hydrogel (Fig. 4G). For instance, in the first triggering cycle, both CS_VEGF_ and CS_PDGF_ were added to the release media at a 0.5:1 mole ratio to aptamers on d3 and d4, respectively. With this ratio, CS_VEGF_ triggered an on-demand VEGF release of 305.9±164.8 pg/ml on d4, while CS_PDGF_ triggered only 5.570±9.65 pg/ml on d5. In the second triggering cycle, increasing the CS to aptamer mole ratio to 2:1, CS_VEGF_ triggered a VEGF release of 364.2±55.76 pg/ml on d8, and CS_PDGF_ triggered a PDGF release of 54.10±80.84 pg/ml on d9 (Fig. 4F&I). For comparison, the same amounts of both CS were supplemented to the GelMA hydrogels at the same time-points. Unlike the DA hydrogels, GelMA hydrogels released only 9.81±9.37 pg/ml of VEGF on d4 (after the first trigger) and around zero pg/ml of VEGF on d8 (after the second trigger). Similarly, with CS_PDGF_, GelMA hydrogels released 81.17±13.256 pg/ml of PDGF on d5 (after the first trigger) and 48.47±42.157 pg/ml on d9 (after the second trigger) (Fig. 4H). It is important to note that the observed release of PDGF/VEGF from the GelMA hydrogels in each triggering cycle is facilitated by a combination of passive diffusion and polymer degradation, rather than the affinity-based triggering effect observed in DA hydrogels. These results indicate that the CS used in this study effectively regulated the binding functionality of aptamers and induced rapid dissociation of aptamer-GF complexes, thereby triggering VEGF/PDGF release from DA hydrogels. Despite the short GF loading time (1 hr) and the relatively low concentration (50 ng) used in this study, CS_VEGF_ facilitated a significantly higher GF release per trigger and similar GF retention percentages compared to other studies using aptamers-modified hydrogels for triggered GF release (Fig. 4B & Table S2, Supporting Information). Higher GF release could potentially be achieved with increased GF loading times and concentrations (Table S2, Supporting Information).

CS-based GF triggering exhibited a mole ratio-dependent behavior, where higher CS to aptamer mole ratios resulted in a greater amount of GF release within 24 hr of triggering, consistent with literature findings.[43] However, with same mole ratios and triggering time, a higher amount of VEGF release was observed from DA hydrogels compared to PDGF. Apart from the triggered GF release behavior, upon triggering DA hydrogels with CS_VEGF_, although the hydrogels showed a maximum VEGF release within 24 hr of triggering, ELISA data indicated that DA hydrogels took almost 2-3 days to revert the GF release to baseline levels. This trend was more prevalent in daily VEGF release profile of DA hydrogels compared to PDGF release. For instance, in each triggering cycle, VEGF levels peak on the triggering day, but DA hydrogels continued to release significant amounts of VEGF even on the second day of triggering (63.49±23.14 pg/ml VEGF on d5 and 143.17±61.99 pg/ml VEGF on d9) (Fig. 4G). The dissociation constant (Kd) of an aptamer is a critical parameter that measures the affinity between the aptamer and its target molecule. The Kd value represents the concentration of the target at which half of the aptamer molecules are bound to their target. A lower Kd value indicates higher affinity and specificity, meaning the aptamer binds strongly to its target even at low concentrations. Conversely, a higher Kd value suggests lower affinity, requiring a higher concentration of the target for effective binding. Therefore, the observed lower amount of triggered PDGF release could be attributed to the higher Kd value of Aptamer_PDGF_ (109 nM)[22], whereas the Aptamer_VEGF_ (370 pmol)[31,32] with a lower Kd could release almost similar amounts of VEGF even with different mole ratios. By modulating the aptamer’s binding affinity and mole ratio, improved GF triggered release can thus be achieved.

DA hydrogels displayed a significantly lower burst release of both GFs compared to GelMA hydrogels on d1 (Fig. S4&5, Supporting Information). Specifically, on d1, DA hydrogels released 260.78±39.40 pg/ml (CV=15.11%) of VEGF and 26.99±23.82 pg/ml (CV=88.25%) of PDGF, whereas GelMA hydrogels released 356.23±46.48 pg/ml (CV=13.05%) of VEGF and 720.75±100.62 pg/ml (CV=13.96%) of PDGF. Notably, while DA hydrogels showed a marked reduction in non-specific GF release after d1, GelMA hydrogels continued to release unbound or freely diffusible GFs, particularly PDGF, whose release was associated with passive diffusion and ongoing polymer degradation up to d5. This observation underscores the enhanced control over initial burst release and the effective dual GF sequestration achieved with DA hydrogels.

The cumulative VEGF release data demonstrates that DA hydrogels achieved a significantly higher release compared to GelMA hydrogels. Specifically, DA hydrogels underwent two cycles of triggered VEGF release from the same samples, yielding a total of 1246.75±93.61 pg/ml (CV=7.51%) on d10, whereas GelMA hydrogels, which predominantly released VEGF in a burst on d1, released only 525.14±69.43 pg/ml (CV=13.22%) by d10 (Fig. 4D). In contrast, DA hydrogels exhibited a lower overall cumulative PDGF release (156.28±153.54 pg/ml, CV=98.25%) compared to GelMA hydrogels (1722.75±336.18 pg/ml, CV=19.51%) on d10 (Fig. 4E). GelMA hydrogels released most of their PDGF in a burst on d1, reflecting non-specific release behavior. The reduced PDGF release from DA hydrogels is likely due to the higher Kd value of Aptamer_PDGF_, suggesting that using aptamers with higher affinity could address this limitation. Nevertheless, the cumulative VEGF release data confirms the proof-of-concept for DA hydrogels, highlighting their capability to achieve substantial programmed GF release. Upon triggering, DA hydrogels release only the GFs, with minimal non-specific release in the absence of an external trigger (CS). This indicates that DA hydrogels can rapidly sequester multiple GFs within an hour and achieve on-demand, programmable release profiles under physiologically relevant conditions by optimizing the mole ratios of the respective CS.

It is also important to note that GelMA hydrogels cumulatively released approximately 525 pg of VEGF and 1723 pg of PDGF, which represents only a small fraction of the 50 ng initially loaded per sample. Analysis of the residual free VEGF in the loading solution indicated that about 28% of VEGF was incorporated into the samples. While this might suggest that a significant portion of the loaded VEGF remains within the GelMA hydrogels after a 10-day release period, this is unlikely given the minimal staining for both VEGF and PDGF in the line micropatterns shown in Fig. 3A. If GelMA hydrogels had strong binding properties sufficient to retain most of the VEGF over the entire period, one would expect negligible impact of GF binding to the aptamers, leading to only minor or undetectable differences in staining. The reason for this discrepancy remains unclear; however, given that GF sequestration and release experiments were conducted separately, variations between individual samples could not be entirely ruled out.

The present GF loading method decouples the loading process from hydrogel fabrication, allowing for flexible GF loading cycles post-synthesis. This approach accommodates the evolving needs of engineered or host tissue by using biocompatible, mild crosslinking processes, ensuring that GF loading and release occur under physiological conditions. Utilizing GelMA as the polymeric backbone for DA hydrogels, which is known for its biocompatibility and suitability for biofabrication,[44] enhances the potential of DA hydrogels for tissue engineering applications. While GelMA degradation could limit GF release in longer culture periods, this can be addressed by modulating polymer concentrations, methacrylation degree, and crosslinking density. An optimal balance of these parameters is essential for maximal aptamer incorporation and matching the hydrogel’s mechanical properties with the desired tissue characteristics. We demonstrated the compatibility of DA hydrogels with biofabrication techniques, such as photo-patterning, and envision that advanced techniques like 3D-bioprinting and two-photon photo-patterning can provide higher-resolution spatial patterns of multiple aptamers, allowing for a more detailed investigation of the effects of spatially-defined dynamic GF presentation on cell behavior.

Conventional approaches often require supraphysiological concentrations of GFs, which can be harmful *in vivo*, potentially leading to implant failure and poor host-implant integration. Physiological levels of VEGFA in healthy humans range from 1-3 pM in blood plasma and tissue interstitial fluid, and increase in response to hypoxia.[45] Similarly, PDGF concentration in human blood serum are 17.5±3.1 ng/ml under normal conditions, rising significantly during injury or wound healing.[3] Current dual or multiple GF delivery systems use concentrations ranging from 100 ng/ml up to 20 µg of VEGF/PDGF per scaffold for *in vitro* and *in vivo* analysis,[12,46,47] relying on non-specific sustained or sequential release from microsphere or nanofiber polymer degradation.[46,47] In contrast, our work employs physiologically relevant GF concentrations, reducing the risk of off-target effects associated with multiple GF delivery.

Aptamers offer significant potential for therapeutic applications *in vivo* despite their vulnerability to nuclease-mediated degradation, which results in a brief half-life in the bloodstream.[25,48] Various nuclease-resistant modifications, such as locked nucleic acids, phosphorothioate backbones, Spiegelmers, and g-quadruplex structures, can markedly extend aptamer stability.[25] Despite enhanced nuclease resistance, aptamer’s low molecular mass makes them prone to rapid renal filtration. However, polymer conjugation has proven effective in reducing renal clearance rates. For instance, PEG-conjugated aptamers can achieve circulating half-lives of up to one day following intravenous administration in mice, compared to 5-10 min for unconjugated aptamers.[49] The VEGF-binding RNA-based aptamer Pegaptanib sodium (Macugen, Eyetech/Pfizer) used for treating macular degeneration incorporates both a 40 kDa PEG conjugation and 2’-O-methyl modifications, resulting in an *in vivo* half-life of 10 (±4) days.[50] These advancements suggest that our developed aptamer-based platform can be readily adapted for *in vivo* and clinical applications.

In native ECM, GFs becomes stabilized and protected after binding to various ECM components, acting as reservoirs that releases multiple GFs as needed. Similarly, we have previously shown that aptamer-tethered VEGF within a 3D hydrogel microenvironment elicits selective cell responses compared to GelMA regions for at least ten days of culture, [24,39] suggesting long-term stability and higher bioavailability than freely diffusible VEGF. When spatially patterned using photo-patterning or 3D-bioprinting, aptamer-tethered VEGF induces superior cell responses, forming longer and luminal vascular networks compared to plain GelMA regions.[39,44] This is due to the higher local concentrations of aptamer-tethered GFs near the cell surface, leading to more sustained and potent signalling. Aptamer-tethering preserves the GF binding domains and protects GFs from proteolytic degradation, providing a stable and prolonged signal compared to freely diffusible GFs, which can be rapidly diluted or degraded in the ECM. Therefore, we envision that this platform, compatible with physiological conditions, can also be used to dynamically present multiple GFs to the cells with 4D control, mimicking native ECM characteristics.

One key advantage of this platform is its ability to trigger the release of two GFs without any leakage or non-specific release, achievable in repeatable cycles at physiological temperature. Mimicking the native ECM’s characteristics as a reservoir for multiple GFs, this platform offers the flexibility to use the same hydrogel matrix as GF reserves or depots, delivering differential GF release amounts at each triggering cycle. State-of-the-art GF delivery approaches face challenges in loading multiple GFs and controlling their individual release profiles. This platform can be leveraged for drug delivery or other biomedical applications, potentially replacing single-use hydrogel implants that require re-implantation after one complete release cycle. Owing to aptamer’s high binding specificities, multiple aptamers can be incorporated within the same hydrogel, allowing for different release kinetics for various GFs. This flexible control over individual GF release kinetics opens new avenues for “intelligently” controlled GF delivery, enabling adjustment of individual release profiles according to the spatiotemporally changing requirements of growing engineered tissue.

## 4. Conclusions

In conclusion, we have developed an “intelligent” biomaterial platform that enables spatial control of the bioavailability of two angiogenic GFs (VEGF & PDGF-BB). Utilizing aptamer affinity interactions with their target GFs and corresponding CS within a biofabrication-friendly polymeric matrix (GelMA), we achieved on-demand, triggered release of individual GFs in multiple cycles and in a time-controlled manner, establishing a “programmable” behavior for tissue engineering application. The platform exhibits several outstanding features: (i) Stable incorporation of dual aptamers, each specific for VEGF and PDGF, within the polymeric matrix through chemical modification; (ii) Functional aptamer-CS molecular recognition in a 3D microenvironment with long-term stability, lasting at least 15 days at physiological temperature; (iii) Rapid and spatially localized dual GF sequestration from the culture medium, confirming high spatial selectivity for individual GFs; (iv) On-demand, triggered GF release via CS hybridization in multiple cycles, allowing differential amounts of individual GFs to be released from the same hydrogels at different time-points. This innovative ability to selectively sequester and release multiple GFs from hydrogels, incubated in a single medium pool, opens up new avenues for mimicking dynamic GF presentation in a 3D matrix, similar to native ECM. Additionally, the platform demonstrated significantly lower burst releases of both GFs compared to control GelMA hydrogels, with no leakage or non-specific release. This flexible control over individual GFs release kinetics offers new possibilities for “intelligently” controlled GF delivery, where release profiles can be adjusted according to the spatiotemporally changing requirements of growing engineered vasculature or tissue.

## CRediT author statement

**Deepti Rana:** Conceptualization, Methodology, Investigation, Formal analysis, Writing-original draft, Writing - review & editing. **Jeroen Rouwkema:** Conceptualization, Writing-review & editing, Supervision, Project administration, Funding acquisition.

## Acknowledgements

This work was supported by the European Research Council (ERC) under the ERC Proof of Concept Program (No. 101062032, BioTisSeal). Illustrations for the manuscript were created with BioRender.com.

## Supplementary Information

Supplementary data to this article can be found online.

