## Supplementary Information for "Spatiotemporally Programmed Release of Aptamer Tethered Dual Angiogenic Growth Factors"

#### Supporting Information

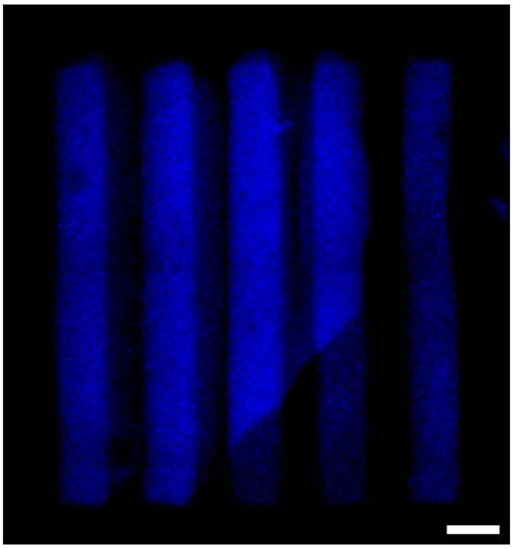

**Figure S1. The proof-of-concept line micropattern designs.** The fluorescence microscopic stitched image of the micropattern fabricated via two-step photocrosslinking method. The blue fluorescence microbeads were mixed with GelMA pre-polymer for the 1st crosslinking cycle followed by plain GelMA for 2nd crosslinking cycle. The same micropattern designs were fabricated using DA and GelMA hydrogels for GF staining experiments. The scale bar is 500 µm.

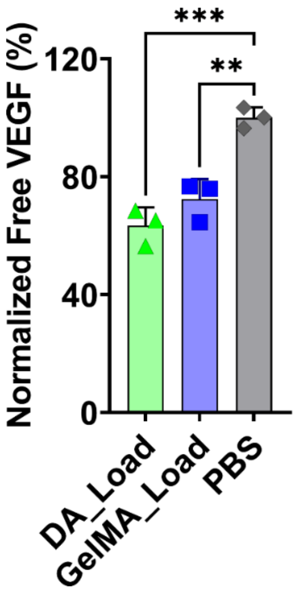


**Figure S2. The free VEGF concentration present in the loading solution after DA and GelMA hydrogel incubation for 1hr.** To measure the ELISA sensitivity for free VEGF molecule detection, same VEGF loading concentration was added into the PBS, followed by 1hr of incubation at 37 °C. The initial VEGF loading concentration for all samples is 50 ng/ml. The data is normalized with control PBS samples and plotted on right-y-axis. The quantification was performed using ELISA assay with three experimental replicates (n=3). The data is represented as mean ± S.D. The statistical significance was calculated using unpaired t-test with Welch's correction where alpha was fixed at 0.05 with ***p=0.0004, and **p=0.0018.

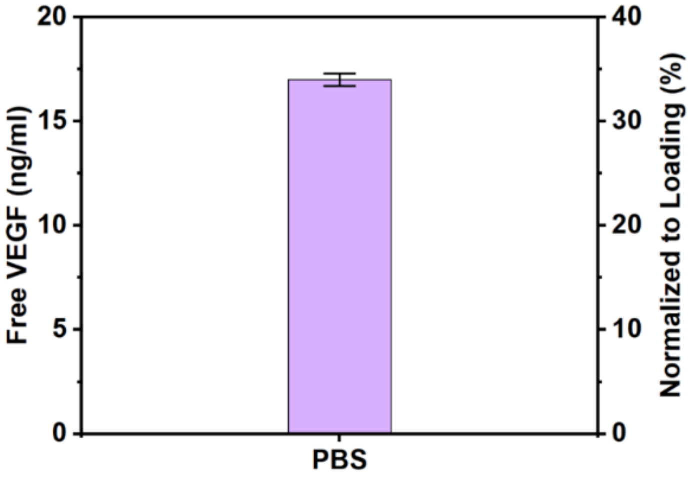


**Figure S3. Actual free VEGF concentration present in the PBS after 24hr incubation at 37 °C.** The PBS samples were loaded with 50 ng/ml of VEGF. The data presents actual free VEGF concentration detected using ELISA assay (left-y-axis) and the detected concentration normalized to the loading concentration (right-y-axis). The graph indicates the difference in ELISA sensitivity in free VEGF detection in the PBS compared to the VEGF loading concentration. The quantification was performed using ELISA assay with three experimental replicates (n=3). The data is represented as mean ± S.D.

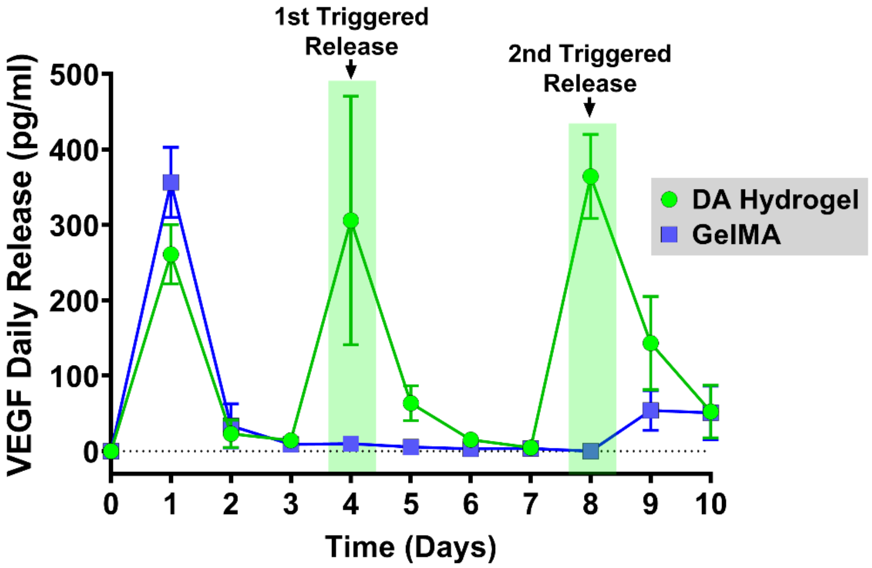


**Figure S4. The daily VEGF release from DA and GelMA hydrogels till d10.** To trigger VEGF release on d4 and d8, the $CS_{VEGF}$ was supplemented to DA hydrogels on d3 and d7, respectively.

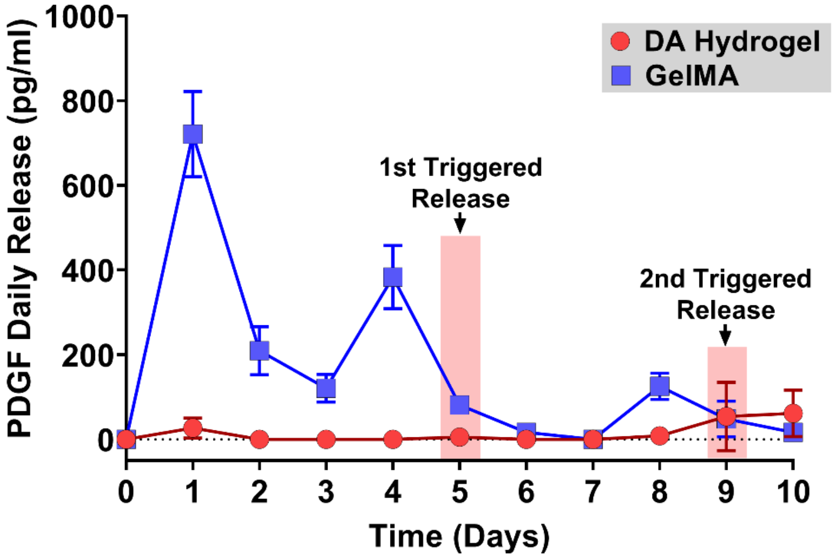


**Figure S5. The daily PDGF release from DA and GelMA hydrogels till d10.** To trigger PDGF release on d5 and d9, the $CS_{PDGF}$ was supplemented to DA hydrogels on d4 and d8, respectively.

### Supporting Tables & Video

**Table S1. Full sequences and other characteristics of the aptamers used in this study.** Tm denotes the melting temperature (50 mM NaCl), MW is molecular weight and N signifies the number of nucleotides.

| Aptamer | Sequence (5'→ 3') | Tm | MW | N |
|---|---|---|---|---|
| **$Aptamer_{VEGF}$** | /5Acryd/CGA TCG TAT CAG TCC ACA AGC CCG TCT TCC AGA CAA GAG TGC AGG GC | 70.8 °C | 14665.6 | 47 |
| **$CS_{VEGF}$** | CGC CCT GCA CTC TTG TCT GGA AGA CGG GCT TGT GGA CTG ATA CGA TCG | 71.3 °C | 14791.6 | 48 |
| **$CS_{F-VEGF}$** | /5Alexa488N/CGC CCT GCA CTC TTG TCT GGA AGA CGG GCT TGT GGA CTG ATA CGA TCG | 71.3 °C | 15487.2 | 48 |
| **$Aptamer_{PDGF}$** | /5Acryd/GC GAT ACT CCA CAG GCT ACG GCA CGT AGA GCA TCA CCA TGA TCC CA | 70.8 °C | 14305.4 | 46 |
| **$CS_{PDGF}$** | TGG GAT CAT GGT GAT GCT CTA CGT GCC GTA GCC TGT GGA GTA TCG C | 70.8 °C | 14244.2 | 46 |
| **$CS_{F-PDGF}$** | /5Alex647N/TG GGA TCA TGG TGA TGC TCT ACG TGC CGT AGC CTG TGG AGT ATC GC | 70.8 °C | 15264.4 | 46 |

**Table S2. Comparative data on GF retention and release properties of aptamer-functionalized hydrogels.** This table compiles the data used to plot the factors influencing GF retention and release properties in various aptamer-functionalized hydrogels, as reported in the literature. The data from the current study were compared with previous studies [1–4] and visualized in a radar chart (Fig. 4B) using Origin software. Values that were unavailable or not applicable are listed as zero. For consistency and ease of comparison, only the release data for the first GF (VEGF or PDGF) from studies reporting dual GF release were included.

| References | GF Release/ Trigger (pg) | GF Retention (%) | CS/Aptamer Mole Ratio | Aptamer/GF Mole Ratio | GF Loading Conc. (ng) | Triggering Time (hr) | GF Loading Time (hr) |
|---|---|---|---|---|---|---|---|
| This study | 364.188 | 37 | 2 | 2000 | 50 | 24 | 1 |
| [1] | 25.481 | 46 | 1 | 10000 | 10 | 24 | 1 |
| [2] | 560 | 0 | 625 | 1500 | 4 | 1 | 0 |
| [3] | 1125 | 41 | 1 | 0.4 | 7500 | 1 | 3.5 |
| [4] | 1500 | 0 | 1.5 | 200 | 15 | 1.5 | 12 |

**Video S1. 3D projection of confocal z-stacks displaying the interface region of micropattern decorated with dual aptamers, specifically Aptamer$_{VEGF}$ and Aptamer$_{PDGF}$.** These micropatterns were incubated for 24hrs at 37 °C with fluorescently labelled CS$_{VEGF}$ (green) and CS$_{PDGF}$ (red). The 3D projection confirms the spatially localized and homogenous sequestration of the aptamer-specific CS throughout the entire micropattern thickness (z=240 μm).

**Video S2. 3D projection of confocal z-stacks of VEGF (red) immunostained dual aptamer patterned line-micropatterns.** Maximum VEGF localization is restricted to the patterned aptamer regions (line width=500 μm; z=240 μm). The blue color represents fluorescent microbeads pre-mixed with the GelMA pre-polymer, indicating the GelMA region (line width=500 μm).

**Video S3. 3D projection of confocal z-stacks of PDGF (green) immunostained dual aptamer patterned line-micropatterns.** Maximum PDGF localization is restricted to the patterned aptamer regions (line width=500 μm; z=240 μm). The blue color represents fluorescent microbeads pre-mixed with the GelMA pre-polymer, indicating the GelMA region (line width=500 μm).

**Video S4. 3D projection of confocal z-stacks of composite VEGF and PDGF immunostained dual aptamer patterned line-micropattern (Video S2 & S3).** The blue color represents fluorescent microbeads pre-mixed with the GelMA pre-polymer, indicating the GelMA region (line width=500 μm).
